# Automethylation of PRC2 fine-tunes its catalytic activity on chromatin

**DOI:** 10.1101/349449

**Authors:** Chul-Hwan Lee, Jeffrey Granat, Jia-Ray Yu, Gary LeRoy, James Stafford, Danny Reinberg

## Abstract

The catalytic activity of PRC2 is central to maintain transcriptional repression by H3K27me3-decorated facultative heterochromatin in mammalian cells. To date, multiple factors have been reported to regulate PRC2 activity. Here, we demonstrate that PRC2 methylates itself on EZH1/2 and SUZ12 subunits, with EZH1/2-K514 being the major automethylation site in cells. The functional studies of automethylation on EZH2 indicate automethylation as a self-activating mechanism for PRC2 in the absence of stimulatory cofactors like AEBP2. Together, our study reveals PRC2 automethylation as a novel regulatory mechanism of PRC2 activity on chromatin.

## Introduction

Once established, transcriptional repression of developmentally silenced genes is maintained by the Polycomb Group (PcG) proteins. The enzymatic complex at the forefront of PcG-mediated gene repression is Polycomb Repressive Complex 2 (PRC2). PRC2 consists of 4 subunits, EZH1/2 (the catalytic subunit), EED, SUZ12, and RBAP46/48, and catalyzes H3K27 mono/di/tri-methylation (H3K27me1/2/3). EZH1 and EZH2 are interchangeable subunits of PRC2, with PRC2-EZH1 being less catalytically active. Upon the catalysis of H3K27 methylation by PRC2-EZH1/2, EED recognizes H3K27me3, the terminal enzymatic product, resulting in allosteric activation of the complex (Margueron et al. 2009). This positive feedback loop generates large spans of H3K27me3 domains (Lee et al.2018b; (Margueron et al. 2009); Oksuz et al. 2018), a hallmark of facultative heterochromatin and transcriptional repression. Failure to maintain these domains contributes to pathogenic conditions including developmental diseases and cancers (Wassef and Margueron 2017). Therefore, it is critical to comprehensively understand the full scope of mechanisms regulating PRC2 catalytic activity. To date, several layers of regulation have been demonstrated, including the nucleosome density on chromatin, PRC2 accessory factors, PRC2 allosteric activators, and mRNAs as well as long non-coding RNAs (Holoch and Margueron 2017). Recent advances further highlight the crucial roles of PRC2 accessory factors such as AEBP2, JARID2, EPOP, LCOR/LCORL, and Polycomb-like (PCL) proteins in regulating the activity and/or recruitment of PRC2 (Li et al. 2017; Lee et al. 2018a; Oksuz et al. 2018; Liefke et al. 2016; Beringer et al. 2016; Son et al. 2013; Sanulli et al. 2015; Conway et al. 2018).

In addition to H3K27, non-histone substrates of PRC2 have also been reported as PRC2 regulators. Most notably, JARID2, a PRC2-interacting protein, was shown to be methylated at lysine 116 (JARID2-K116me) by PRC2. JARID2-K116me3 then elicits a positive feedback response on PRC2 by allosterically stimulating its catalytic activity (Sanulli et al. 2015). Additionally, Elongin A was recently shown to be methylated by PRC2 and subsequently regulated the expression of PRC2 target genes (Ardehali et al. 2017). These findings highlight the important role of non-histone substrates in regulating the function of PRC2.

Recent studies employing histone methyltransferase (HMT) assays indicated that PRC2 incorporates the methyl group of radioactive S-adenosylmethionine (SAM) into its own subunits (Sanulli et al. 2015; Wang et al. 2017). Since a number of enzymes coordinate feedback loops through self-modification (e.g. autophosphorylation or automethylation) (Chin et al. 2007; Wassef and Margueron 2017Wang and Wu 2002; Dillon et al. 2013), we speculated that PRC2 automethylation might similarly serve an important role in the regulation of PRC2 HMT activity.

Here, we investigate the functional significance of PRC2 automethylation as a novel regulatory mechanism on the catalytic activity of PRC2. Using reconstituted *in vitro* systems and mass spectrometry, we identified the sites of PRC2 automethylation and further tested their impact on PRC2 activity using both *in vitro* and *in vivo* cellular systems.

## Results and Discussion

### Human PRC2 methylates itself *in vitro*

To investigate the possibility that PRC2 methylates itself, we first employed an *in vitro* methyltransferase assay with the PRC2 complex comprising EZH2, SUZ12, EED, and RBAP48. Interestingly, we found that PRC2 automethylated EZH2 and to a lesser extent SUZ12, but not EED or RBAP48 (Fig. 1A, lanes 1 and 2). We next asked if the addition of AEBP2, a PRC2 cofactor that plays a robust stimulatory role, can augment such effect on EZH2/SUZ12 automethylation. Unexpectedly, the PRC2-AEBP2 complex manifested a lower level of automethylation on EZH2/SUZ12 with a nearly complete loss of SUZ12 methylation (Fig. 1A, lanes 3 and 4). In contrast, the presence of an H3K27me3 peptide allosterically stimulated PRC2 and substantially increased the levels of EZH2/SUZ12 automethylation (Fig. 1A, lanes 5 and 6). Next, we performed a standard histone methyltransferase (HMT) assay for PRC2 using oligonucleosomes as an additional substrate (Fig.1A, lanes 7-12). As expected, PRC2 methylated histone H3 at lysine 27; however, the addition of oligonucleosomes had a minor impact on the automethylation of EZH2 and SUZ12. Interestingly, the PRC2-AEBP2 complex manifested increased HMT activity but similar automethylation activity compared to the PRC2. Together, these results suggest that AEBP2 negatively regulates PRC2 automethylation possibly by a switch of its substrate preference, and/or partially hindering EZH2 automethylation residue(s) (see below). This effect can be counteracted by the presence of oligonucleosomes as AEBP2 nucleosome binding further stimulates PRC2 (Lee et al. 2018a) and/or the methylated nucleosomes may in turn stimulate PRC2 by a previously described feed-forward mechanism (Lee et al.2018b). Similarly, the PRC2-EZH1 complex also manifested EZH1/SUZ12 automethylation (Fig. 1B), and the addition of AEBP2 again diminished SUZ12 automethylation (Fig. 1C). Further, the level of automethylation of PRC2-EZH1 is considerably lower than that of PRC2-EZH2 (Fig. 1C), consistent with previous studies (Lee et al. 2018a; Margueron et al. 2008; Son et al. 2013) showing that PRC2-EZH1 is less catalytically active than PRC2-EZH2.

**Figure 1.**
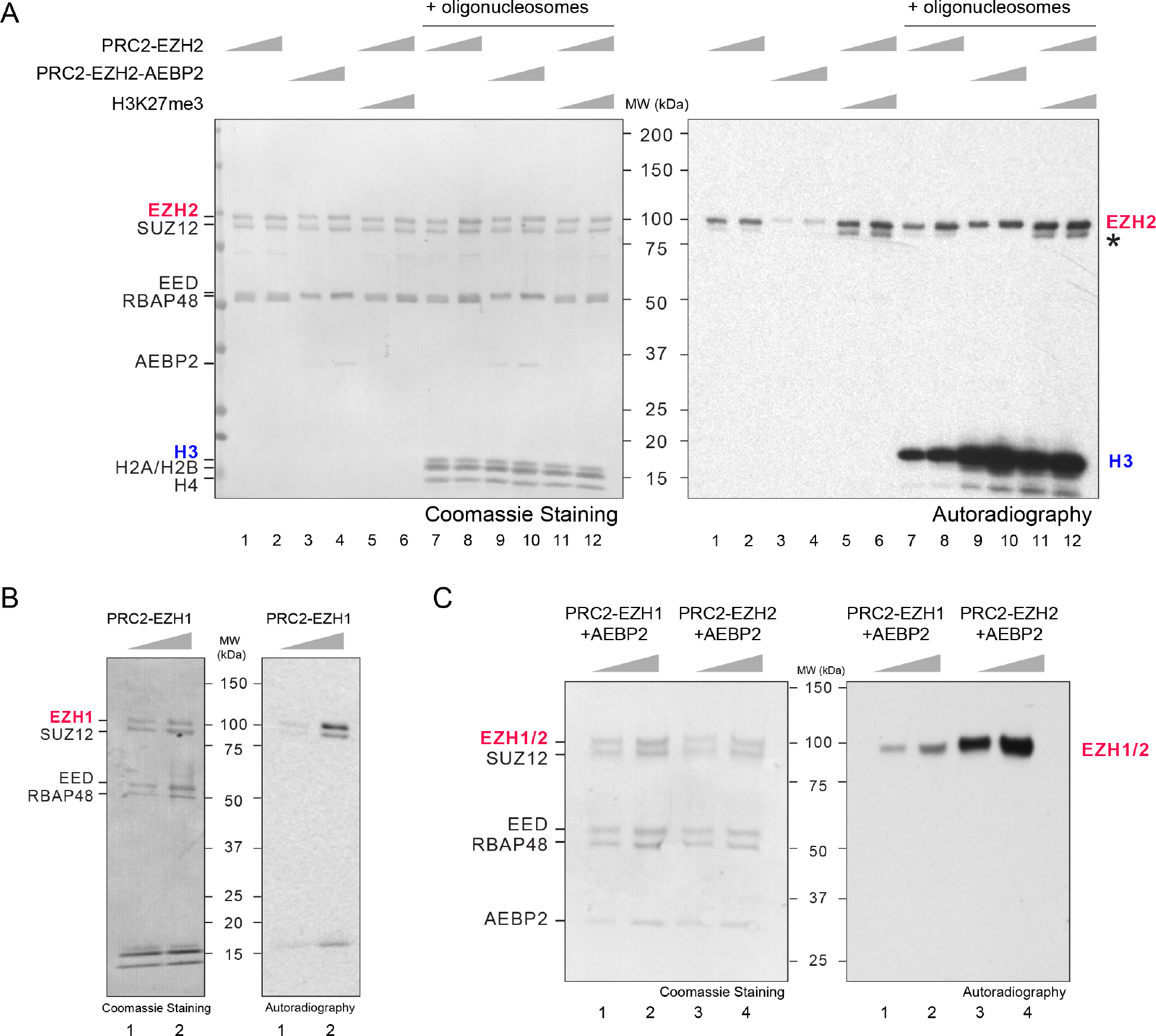
PRC2 subunits are auto-methylated. (A-C) Methyltransferase (MT) assays. The details of MT assay conditions are described in Materials & Methods. (A) MT assays containing PRC2-EZH2, PRC2-EZH2-AEBP2 or PRC2-EZH2 with H3K27me3 peptide (30 or 60 nM) in the absence (lanes 1-6) or presence (lanes 7-12) of oligonucleosomes as substrates (300 nM). Coomassie blue staining of SDS-PAGE gels containing nucleosomes or PRC2 components (*Left* image) was used to visualize the relative concentration of each component present in each reaction. The levels of methylation on EZH2, SUZ12 (asterisk) or histone H3 are shown by autoradiography (*Right* image). (B) MT assays containing PRC2(EZH1) (30 or 60 nM) using oligonucleosomes as substrates (300 nM). Left/Right images are described in Figure 1A. (C) MT assays containing PRC2-EZH1 or PRC2-EZH2 with AEBP2 (30 or 60 nM). Left/Right images are described in Figure 1A.

### Automethylation of EZH1/2 occurs in a key regulatory region between SANT2 and CXC domains

To better understand the functional role of PRC2-EZH1/2 automethylation, we performed mass spectrometry (MS) on *in vitro* automethylated PRC2 to identify the automethylated residues. We identified three methylated lysine residues, all of which reside within an unstructured but highly conserved region of EZH1 and EZH2, including EZH2-K510, −K514, and −K515, corresponding to EZH1-K511, −K515, −K516 (Fig. 2A); specifically, we detected EZH2-K510 monomethylation (K510me1), EZH2-K514 mono-, di- and tri-methylation (K514me1/2/3), and EZH2-K515 di-methylation (K515me2) from our *in vitro* automethylated PRC2 (Figs. 2B-2D). In light of the recent structural studies of PRC2, we realized that these residues reside in the region wherein EZH2 is in close proximity to the H3 tail extending from the nucleosome core (Poepsel et al. 2018). More precisely, the automethylated residues likely contact a region of the H3 tail including amino acid 35-42, which is critical for histone H3 recognition by PRC2 (Yuan et al. 2012). It is also possible that these lysine residues contribute to the interaction between PRC2 and nucleosomes through their electrostatic interactions with nucleosomal DNA (Poepsel et al. 2018). Furthermore, this region of EZH2 lies at the interface between PRC2 and AEBP2 (Ciferri et al. 2012; Poepsel et al. 2018; Kasinath et al. 2018), a well-established regulator of PRC2 catalytic activity. We therefore speculate that EZH1/2 automethylation regulates PRC2 activity by: 1) facilitating the sensing of the H3 tail and/or nucleosomal DNA for efficient catalysis; and 2) modulating the interactions among PRC2, AEBP2 and the nucleosome substrate. Hereon, we focused on PRC2-EZH2, since PRC2-EZH1 and PRC2-EZH2 have the same sites of automethylation.

**Figure 2.**
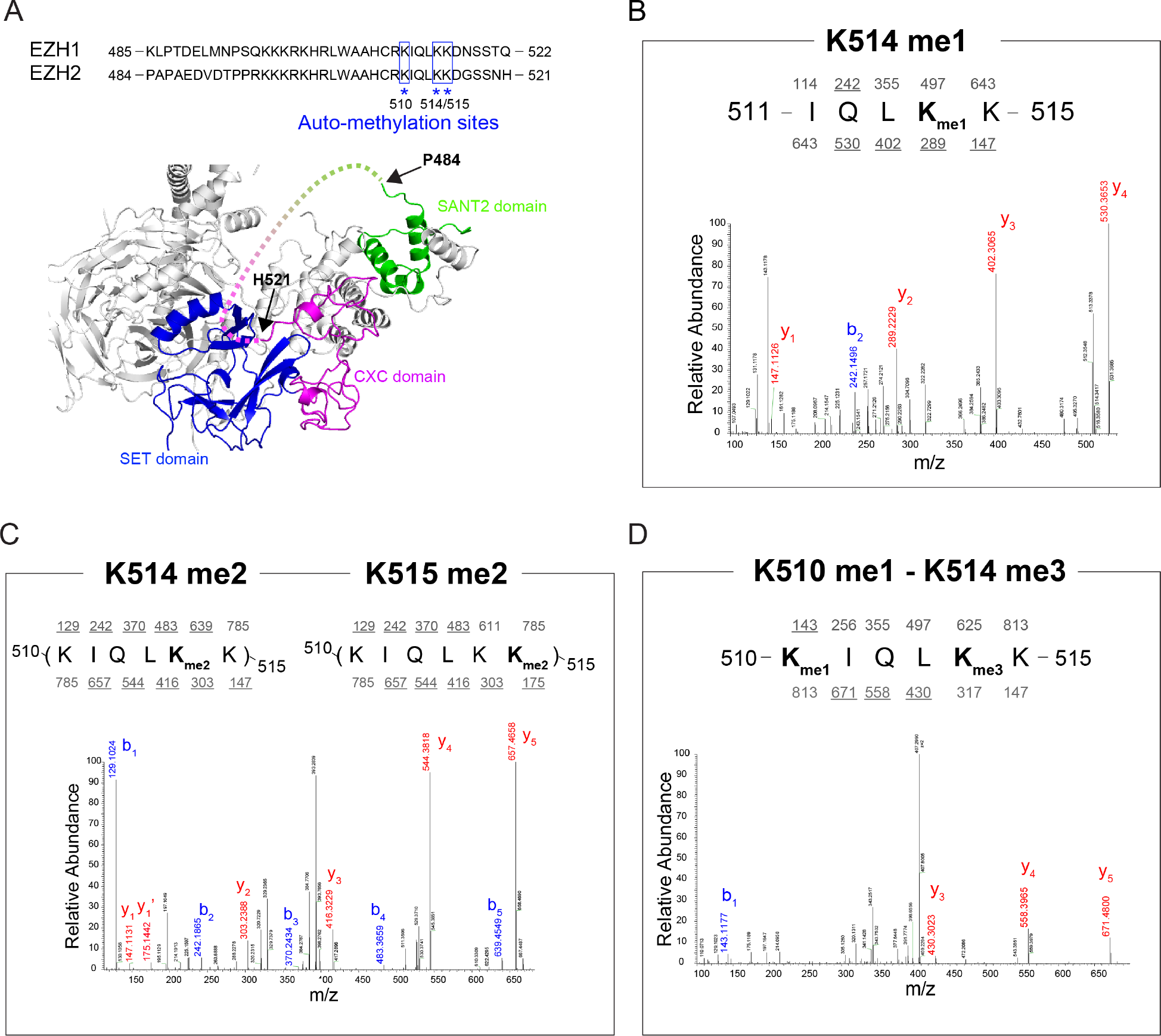
Identification of EZH2 methylation sites on recombinant PRC2 complex. (A) Top, Sequence alignment of human EZH1 and EZH2 between SANT2 and CXC domains. Automethylation sites identified by mass spectrometry (MS) were highlighted by blue square and asterisk. *Bottom*, Human PRC2 structure showing the position of unstructured regulatory loop (dash line) between SANT2 and CXC domains (Modified from PDB: 5HYN). (B-D) MS analyses of automethylated recombinant PRC2 complex. Peptides were generated from a trypsin digestion. MS/MS spectrum showing that EZH2 Lys-514 is monomethylated (B), Lys-514 or 515 is dimethylated (C), and Lys-510 is monomethylated while Lys-514 is trimethylated (D). It should be noted that values above and below the sequence refers to nominal masses for predicted *b* and *y* fragment ions, respectively, with masses underlined indicating the observed fragments.

### Residues automethylated in EZH2 are critical for PRC2 catalytic activity

To test our hypothesis on PRC2 automethylation, we generated EZH2-K514A/K515A and EZH2-K514R/K515R mutants (EZH2-KK514,515AA/RR as EZH2^KKAA/RR^ hereafter) as these two residues were mainly methylated at higher orders (di-and tri-methylation) and K510 was only mono-methylated in relatively low abundance (Fig. 2D and see below). Note that we not only mutated lysine to alanine but also to arginine in order to preserve the positive charge. We first performed an HMT assay using PRC2-EZH2^WT^ or PRC2-EZH2^KKAA/RR^ and demonstrated that mutations of these residues abolished the automethylation of PRC2, reaffirming our previous MS results that EZH2-K514/K515 are major sites of automethylation (Fig. 3A). Importantly, the PRC2-EZH2^KKAA/RR^ mutants showed an impaired HMT activity toward oligonucleosomes, underscoring an important role of PRC2 automethylation on regulating its own activity. The addition of AEBP2 again significantly enhanced the HMT activity of PRC2-EZH2^WT^ with a marginal increase of EZH2 automethylation. However, in the presence of AEBP2, the PRC2-EZH2^KKAA/RR^ complexes still showed a substantial reduction in HMT activity. Although AEBP2 seems to inhibit PRC2 automethylation, these results suggest that PRC2 automethylation and AEBP2 might have non-redundant roles in promoting the HMT activity of PRC2.

**Figure 3.**
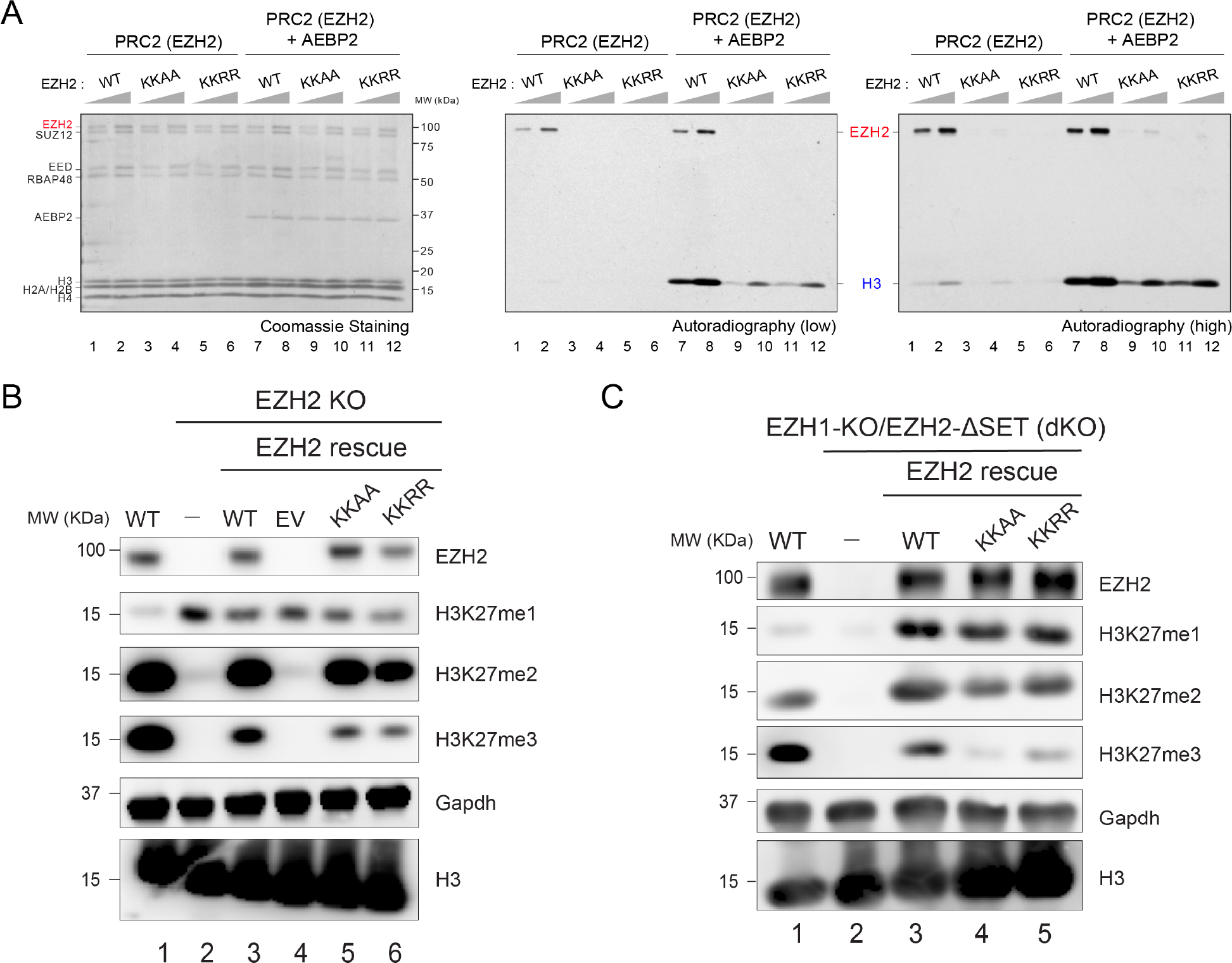
Point mutations of EZH2 methylation sites inhibit PRC2 activity *in vitro* and negatively regulate global H3K27me3 level *in vivo*. (A) Methyltransferase assays using wild-type (WT) PRC2 or mutant PRC2 (30 or 60 nM), as indicated, using oligonucleosomes (300 nM) as substrate. KKAA: EZH2-K514A/K515A, KKRR: EZH2-K514R/K515R. Coomassie blue staining of SDS-PAGE gels containing nucleosomes or PRC2 components (*Left* image) was used to visualize the relative concentration of each component present in each reaction. The levels of methylation on EZH2 or histone H3 are shown by autoradiography (*Middle*/*Right* image). (B and C) Western blot analysis of EZH2, H3K27me1, H3K27me2, H3K27me3 and total histone H3 levels in E14 mESC cells, including WT, EZH2-KO, and EZH2 rescue conditions (B) or B6 mESC cells, including WT, EZH1-KO/EZH2ΔSET (dKO), and EZH2 rescue conditions (C). Gapdh was used as a loading control. mESC, mouse embryonic stem cells. EV, empty vector.

To assess the *in vivo* relevance of these observations, we rescued EZH2 knockout (EZH2-KO) mouse embryonic stem cells (mESCs) with either EZH2^WT^ or EZH2^KKAA/RR^ mutants. mESCs lacking EZH2 showed a nearly complete loss of H3K27me3 (Fig. 3B, lane 2), and cells rescued with EZH2^WT^ showed an almost complete restoration of H3K27me3 levels (Fig. 3B, lane 3). Importantly, mESCs rescued with EZH2^KKAA/RR^ displayed significantly lower H3K27me3 levels relative to EZH2-WT rescued cells (Fig. 3B, lanes 3-6). We performed the same experiment in an EZH1-KO/EZH2ΔSET mESCs, EZH1/2 double knockout (dKO) background and observed an even more profound depletion in H3K27me3 levels when cells were rescued with EZH2^KKAA/RR^ (Fig. 3C). We noted that H3K27me1/2 levels appear to be similar between EZH2^WT^ and EZH2^KKAA/RR^ rescued cells, indicating that EZH2 automethylation is mainly required for higher order H3K27 methylation reminiscent of a previously established role of AEBP2 and PCL1 (PHF1) (Lee et al. 2018a; Sarma et al. 2008). In addition, as the EZH2^KK/AA^ and EZH2^KK/RR^ mutants manifested catalytic deficiencies to a similar degree both *in vitro* and *in vivo*, the positive charge of lysine residues at EZH2-K510, −514, and −515 unlikely plays a role in regulating PRC2 activity.

### EZH1/2-K514 is the major automethylation site in mouse ESCs and 293TREX cells

Having examined the function of EZH2 automethylation *in vitro*, we next examined the existence of these modifications in a cellular context. We took advantage of a previously constructed EED-deficient mESC line wherein a FLAG-tagged EED was ectopically expressed (Lee et al.2018b). PRC2 was purified through FLAG-based affinity purification (Fig. 4A, Left) and then subjected to MS analysis. Importantly, this *in vivo* purified PRC2 also contained automethylation at EZH1/2-K510 and-K514 (Fig. 4B-4D). However, the EZH1/2-K515 and SUZ12 automethylation were not detected in this experiment, possibly due to their relatively low abundance in this cell type. In a similar experiment, a comparable pattern of EZH1/2 automethylation was also observed in 293TREX cells (Fig. 4B). Strikingly and interestingly, when analyzing all the methylated 510-KIQLKK-515 peptides, K514 was mostly methylated and K510 methylation was predominantly observed in the presence of K514me3 *in cis* (Fig. 4B). Together, we conclude that the major site of automethylation in cells is likely EZH1/2-K514, and K514me3 might be a prerequisite for methylation of K510 (Fig. 4). The automethylation on these residues can be inhibited by the presence of AEBP2 (Fig. 1A, lanes 3-4) or possibly other PRC2 cofactors. We propose that automethylation is a self-activating mechanism for PRC2 in a context that is devoid of other cofactors like AEBP2 (Fig. 4E).

**Figure 4.**
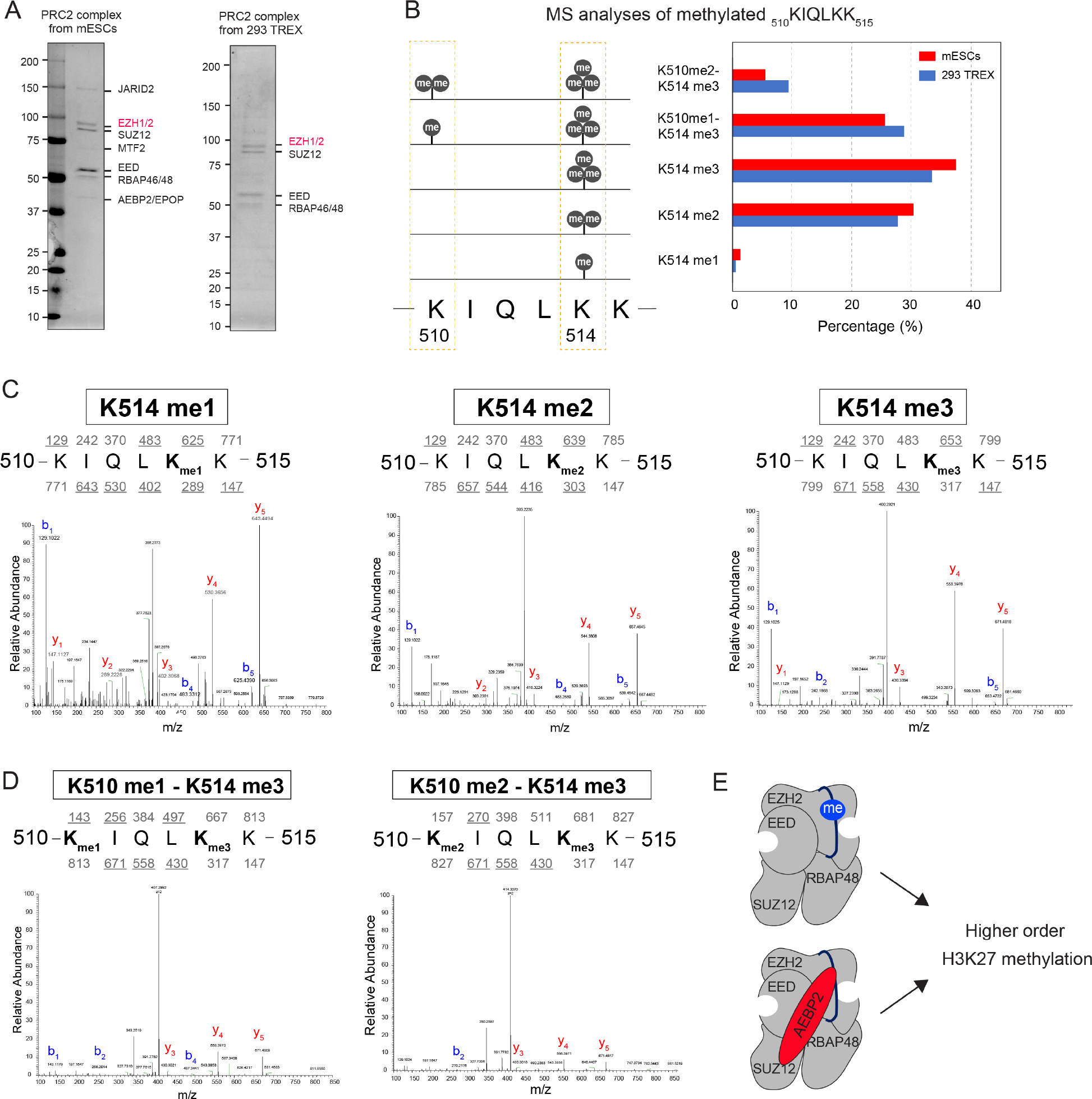
EZH2 K514 is the main methylation site *in vivo*. (A) PRC2 complex was purified from EED FLAG-tagged E14 mESCs (*left*) or 293 TREX cells (*right*). The details of purification are indicated in Materials and Methods. Purified PRC2 complex was shown in Coomassie Blue stained SDS-PAGE gel. (B) Summary of MS analyses of PRC2 complexes purified in panel A. *Left*, illustration of existing methylated peptides (510-KIQLKK-515) in mESCs and 293TREX cells. *Right*, bar graph showing the percentage of each methylation state from the methylated peptide. (C-D) MS/MS spectrum showing that Lys-514 is mono-/di-/tri-methylated (C) and Lys-510 is either monomethylated or dimethylated while Lys-514 is trimethylated (D). It should be noted that values above and below the sequence refers to nominal masses for predicted *b* and *y* fragment ions, respectively, with masses underlined indicating the observed fragments. (E) A working model depicting EZH2 automethylation as a self-activating mechanism for PRC2 while it is not in contact with cofactors like AEBP2.

During the development of our studies, a report from T. Cech’s laboratory was uploaded on June 8 to BioRxiv (doi: https://doi.org/10.1101/343020), showing that EZH2, the catalytic subunit of PRC2 is auto-methylated at specific residues within the EZH2 disordered loop with consequences to its enzymatic activity. While their overall results are consistent with ours, our study differs in the combination of EZH2 automethylation sites examined and their impact on catalysis *in vivo*. Given the overlapping nature of these studies, we chose to share our results as well, as it might prove beneficial to the progress in the field if the findings obtained thus far from both groups on this important topic are shared online.

## Materials and Methods

### Cell culture

Mouse embryonic stem cells (mESCs) were grown in standard ESC medium containing Lif, 1 μM MEK1/2 inhibitor (PD0325901) and 3 μM GSK3 inhibitor (CHIR99021). 293TREX cells were grown in standard DMEM medium supplemented with 10% FBS, 1% non-essential amino acid, 1 mM Na pyruvate, 1% Penicillin/Streptomycin.

### Purification of protein using Baculovirus expression system

To purify human PRC2 complexes, FLAG-tagged-EED, EZH1, EZH2, SUZ12, RBAP48, and Strep-AEBP2 (short isoform) were cloned independently into a baculovirus expression vector, pFASTBac1 (Invitrogen). EZH2 mutant constructs were generated by site-directed mutagenesis and mutations were confirmed by Sanger DNA sequencing. To purify 4 subunit PRC2 complex, all four components (FLAG-tagged-EED, EZH1 or EZH2, SUZ12, and RBAP48) were co-expressed in Sf9 cells by baculovirus infection. To purify 5 subunit PRC2-EZH2 complex, all five components (FLAG-tagged-EED, EZH2, SUZ12, RBAP48 and Strep-AEBP2) were co-expressed in Sf9 cells by baculovirus infection. After 60 hr of infection, Sf9 cells were resuspended in BC150 buffer (25 mM Hepes-NaOH, pH 7.5, 1 mM EDTA, 150 mM NaCl, 10 % glycerol, 0.2 mM DTT, and 0.1 % NP-40) with protease inhibitors (1 mM phenylmethlysulfonyl fluoride (PMSF), 0.1 mM benzamidine, 1.25 mg/ml leupeptin and 0.625 mg/ml pepstatin A) and phosphatase inhibitors (20 mM NaF and 1 mM Na_3_VO_4_). Cells were lysed by sonication (Fisher Sonic Dismembrator model 100), and WT or mutant recombinant PRC2 was tandemly purified through Q sepharose beads (GE healthcare), FLAG-M2 agarose beads (Sigma), and glycerol gradient (15-35%) sedimentation. AEBP2 was also separately purified as described above with exception of using BC350 buffer while purification.

### Nucleosome reconstitution

Recombinant histones were generated as previously described (Yun et al. 2012; Lee et al. 2013). Briefly, each core histone was expressed in Rosetta (DE3) cells (Novagen), extracted from inclusion bodies, and purified by sequential anion and cation chromatography. For refolding recombinant octamers, equal amounts of histones were mixed and dialyzed into refolding buffer (10 mM Tris-HCl, pH 7.5, 2 M NaCl, 1 mM EDTA, and 5 mM β-mercaptoethanol). Octamers were further purified by size exclusion chromatography on a 24-mL Superdex 200 column (GE healthcare) in refolding buffer. Recombinant oligonucleosomes were reconstituted by sequential salt dialysis of octamers and plasmid having 12 repeats 601-nucleosome positioning sequences.

### HMT assay

Standard HMT assays were performed in a total volume of 15 μL containing HMT buffer (50 mM Tris-HCl, pH 8.5, 5 mM MgCl_2_, and 4 mM DTT) with 500 nM of ^3^H-labeled S-Adenosylmethionine (SAM, Perkin Elmer), 10 nM (500 ng) of recombinant oligonucleosomes consisting of 12x repeat nucleosome arrays (300 nM of nucleosome), and recombinant human PRC2 complexes. The reaction mixture was incubated for 60 min at 30 °C and stopped by the addition of 4 μL SDS buffer (0.2 M Tris-HCl, pH 6.8, 20% glycerol, 10% SDS, 10 mM β-mercaptoethanol, and 0.05% Bromophenol blue). A titration of PRC2 (from 5 to 60 nM) was performed under these conditions to optimize the HMT reaction within a linear range, and the yield of each HMT reaction was measured using the following procedures. After HMT reactions, samples were incubated for 5 min at 95 °C and separated on SDS-PAGE gels. The gels were then subjected to Coomassie blue staining for protein visualization or wet transfer of proteins to 0.45 μm PVDF membranes (Millipore). The radioactive signals were detected by exposure on autoradiography films (Denville Scientific).

### Lentiviral production and delivery

WT or mutant EZH2 constructs were subcloned into the pLV-EF1-alpha-IRES-mCherry vector (Clontech) for lentiviral production and delivery. For the production of viral particles, lentiviral vectors containing WT or mutant EZH2 (10 μg) were co-transfected with pcREV (2.5 μg), BH-10 (3 μg), and pVSV-G (5 μg) packaging vectors into 293-FT cells. The virus-containing medium was collected 48 hr after transfection and the target cells were spin-infected. Polybrene was added to the viral medium at a concentration of 8 μg/mL. Infected cells were FACS-sorted for mCherry.

### Purification of PRC2 complex from mESC or 293TREX cells

To purify PRC2 complex from mESC, Flag-tagged EED in EED-deficient background mESCs were used (Lee et al.2018b). Approximately 3×10^8^ cells were cultured and prepared for mESC nuclear extract. Cells were harvested and washed with PBS. Cells were lysed with intact nuclei in TMSD buffer (40 mM Tris-HCl, pH 7.5, 5 mM MgCl_2_, 250 mM sucrose, 1 mM DTT, and 0.02% NP-40) containing protease inhibitors and phosphatase inhibitors, as indicated above. The cell suspension was centrifuged at 800 x g at 4 °C for 10 min. The pellets (nuclei) were resuspended in 20x cell pellet volume of BC400 buffer (20 mM Tris-HCl, pH 7.9, 400 mM KCl, 0.2 mM EDTA, 20% glycerol, 0.5 mM DTT, and 0.02% NP-40) with protease inhibitors and phosphatase inhibitors, and incubated at 4 °C for 1 hr. The cell suspension was centrifuged at 18,000 x g at 4 °C for 20 min. The supernatant was collected as the nuclear extract. The extract was subjected to Flag affinity purification using FLAG-M2 agarose beads. To purify PRC2 complex from 293TREX, Flag-tagged EED was transfected by lentivirus production and delivery. The procedure was described as above.

### Mass spectrometry analyses for PRC2 methylation

Approximately 20 μg (200 nM) of recombinant PRC2-EZH1 or PRC2-EZH2 (FLAG-tagged-EED, EZH1 or EZH2, SUZ12, and RBAP48) was incubated with 1x HMT buffer containing 5 μM of SAM at 30 °C for 2 hrs. After automethylation, the recombinant PRC2 complex was prepared for mass spectrometry analyses. We also isolated the endogenous PRC2 complex from mESCs and 293TREX cells (see above) for mass spectrometry analyses.

For mass spectrometry, approximately 5 μg (30 μl) of purified complex was first denatured with 90 μl of 8 M Guanidine and then reduced with 3 μl of 1M DTT at 60 °C for 30 min. Methionines were then alkylated with 20 mM Iodoacetamide (IAM) at 25 °C overnight. The samples were then digested with trypsin at 37 °C overnight. Following digestion, the peptides were purified by passing the samples through a 10kDa filter followed by 4 washes with 50mM ammonium bicarbonate. Peptides were then concentrated to 20 μl with a speedvac and 1 μl of 10% TFA was added. Samples were then subjected to LC-MS/MS in a Thermo Velos Mass Spectrometer where the MS2s were collected in data dependent acquisition mode. Spectra were then searched with Mascot allowing a static modification of +56 Da for methionines and variable searches for methylated lysines i.e: me1 = +14 Da, Me2 = +28 Da and Me3 = +42 Da, respectively.

## Acknowledgement

We thank Drs. L. Vales for critical reading of the manuscript as well as past and current Reinberg’s lab member for critical comments and discussion; D. Hernandez for technical assistance. We also thank the NYU Flow Cytometry Core (grant: NIH/NCI P30CA016087) for cell purification. We thank H. Zheng from the Biological Mass Spectrometry Facility at Robert Wood Johnson Medical School and Rutgers for Mass spectrometry analyses. The work in D.R.’s lab is supported by NIH (R01CA199652) and the Howard Hughes Medical Institute (HHMI). J-R.Y. is supported by the American Cancer Society (PF-17-035-01). J.G is supported by The Vilcek Scholarship. J.M.S. was a Simons Foundation’s Junior Fellow and is now supported from an NIH (K99AA024837) grant.

## Author contributions

C-H.L., J.G., J-R.Y., and D.R. conceptualized and designed the study. C-H.L., J.G., J-R.Y., G.L. and J.M.S. conducted the experiments. C-H.L., J.G., J-R.Y. wrote the manuscript with assistance from D.R.

